# Separating biological variance from noise by applying EM algorithm to modified General Linear Model

**DOI:** 10.1101/2024.09.29.615661

**Authors:** Tien-Wen Lee

## Abstract

**Introduction:** The General Linear Model (GLM) has been widely used in research, where error term has been treated as noise. However, compelling evidence suggests that in biological systems, the target variables may possess their innate variances.

**Methods:** A modified GLM was proposed to explicitly model biological variance and non-biological noise. Employing the Expectation and Maximization (EM) scheme can distinguish biological variance from noise, termed EMSEV (EM for Separating Variances). The performance of EMSEV was evaluated by varying noise levels, dimensions of the design matrix, and covariance structures of the target variables.

**Results:** The deviation between EMSEV outputs and the pre-defined distribution parameters increased with noise level. With a proper initial guess, when the noise magnitude and the variance of the target variables were similar, there were deviations of 3% and 10–16% in the estimated mean and covariance of the target variables, respectively, along with a 1.7% deviation in noise estimation.

**Conclusion:** EMSEV appears promising for distinguishing signal variance from noise in biological systems. The potential applications and implications in biological science and statistical inference are discussed.

## Introduction

The General Linear Model (GLM), typically formulated as Y = Λ*β + ε, has been extensively utilized in research. Y and Λ represent the dependent and independent variables, while β and ε denote the unknown mean effects and the unexplained errors, respectively. Despite its concise formulation, the GLM has spawned numerous variations to embrace diverse analytical needs. One such extension is the Generalized Linear Model (McCullagh, 2019), which accommodates response variables with the error distributions that deviate from the Gaussian. Another important variant is the Mixed-Effects Model (McCulloch et al., 2001), which integrates both fixed and random effects, making it particularly suitable for hierarchical or nested data structures. Additionally, the Multivariate Generalized Linear Model extends the GLM framework to handle multiple dependent variables simultaneously. These adaptations have substantially broadened the applicability of GLM techniques across various statistical modeling scenarios, thereby enhancing the ability to analyze complex data structure with greater accuracy and precision.

Notably, the treatment of the error term ε reveals a significant distinction between biological and non-biological systems. In engineering contexts (non-biological), it is conventional to attribute all unaccounted variance to noise. Consequently, under the assumption that ε is independently and identically distributed (i.i.d.), the variance of β can be derived as var(ε)x(Λ^H^Λ)^-1^. However, in biological systems, compelling evidence suggests that biopsychological indicators (e.g., β) possess intrinsic variances that warrant separate consideration. In psychophysics, the principle of scale invariance relating the mean and variance is encapsulated by Weber’s law (e.g., Fig.2 in (Chater and Brown, 2008)). Neurophysiological research has demonstrated that stimulus onset can induce a reduction in neural variability (Churchland et al., 2010). These (and many other) findings underscore that ε in biological system is not merely “noise” but encompasses components of physiological significance that have been historically overlooked in the GLM framework.

This investigation commenced with an explicit modeling of the variance components in the GLM, expressed as Y = Λ* β + ε, where β ∼ N(m,Ф) and ε ∼ N(0,Ψ). In this formulation, m and Φ represent the mean vector and covariance matrix of β, respectively, while Ψ denotes the covariance matrix of ε, under the assumption of multivariate normal distributions and i.i.d. This approach diverges from the conventional GLM by explicitly accounting for the variability in both the coefficients (β) and the error term (ε). To estimate the parameters of this modified GLM, the Expectation-Maximization (EM) algorithm was employed. The EM algorithm, an iterative method for finding maximum likelihood estimates, was particularly suitable due to its ability to handle the multiple hidden variables introduced in this model, named EMSEV for convenience hereafter (**EM** for **SE**parating **V**ariances).

Subsequently, simulations were conducted to examine the validity of EMSEV using a public dataset. A strategy of “sliding design matrix” was utilized that leveraged existing psychological or physiological experimental structures as a basis, thus eliminating the need for extensive repetitive experiments. The performance of EMSEV was examined by testing it across various degrees of noise, titrating the ratios between the variances of ε and β from 0.1 to 10. Additionally, different covariance structures of β were explored. The contexts in which this modified GLM finds application are also discussed, providing insights into its potential utility across diverse research scenarios.

## Materials and Methods

### Mathematical derivation for solving EMSEV (refer to **Supplementary Material** for detail)

The modified GLM is formulated as below: Yn = Λ*β + ε, where β ∼ N(m,Ф), ε ∼ N(0,Ψ), n ∈ 1∼N (N samples). Let □ denote parameter space {m,Ф,Ψ}.

**E step:** calculating the posterior distribution of β at data point n (□ is obtained from previous M step)

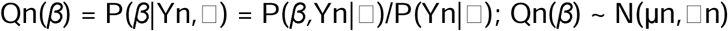

The denominator P(Yn,□) is irrelevant to β, so just focus on the nominator:

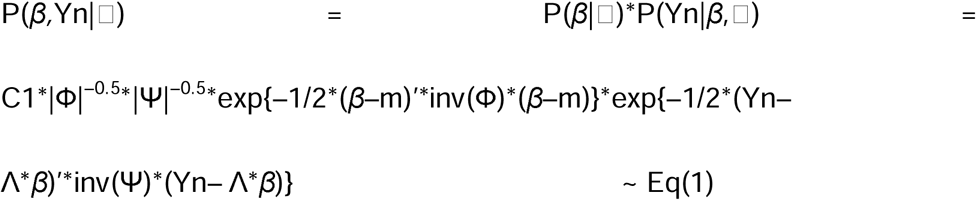

Since the product of Gaussian probability density functions retains a Gaussian distribution, the focus shifts to the exponential component, and rewrite Eq(1) to obtain:

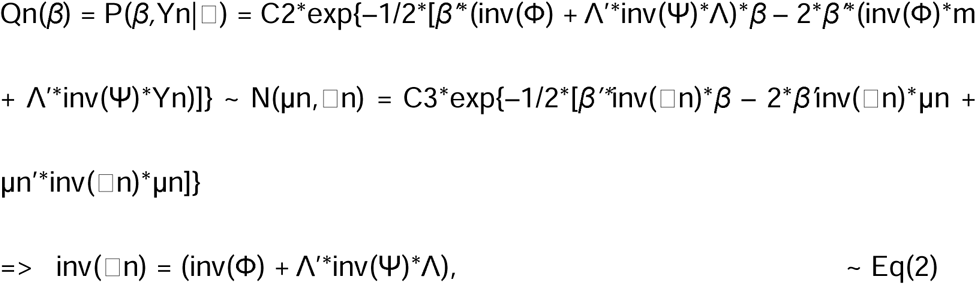

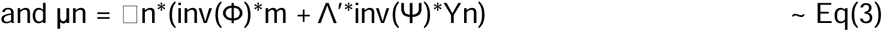

The distribution of Qn(β) is thus derived. For a fixed design matrix Λ, □n = □ since it’s irrelevant to sample Yn. □ is used hereafter. Additionally, it is evident that

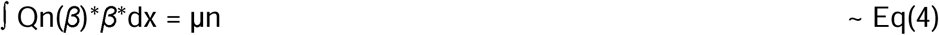

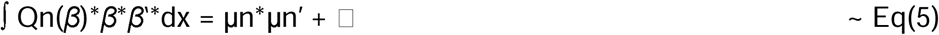

The lower bound of the log-likelihood of the entire dataset is ∑ log(P(Yn|□)) =

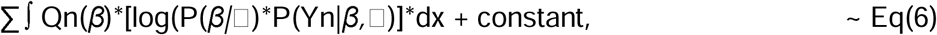

where ∑ denotes summation over N samples (the same applies below).

**M step:** taking derivative of Eq(6) with respect to □

For now, focus on ∫ Qn(β)*□log(P(β|□)*P(Yn|β,□)]*dx for a particular sample n, i.e., taking expectation of log(P(β|□)*P(Yn|β,□) over Qn(β)

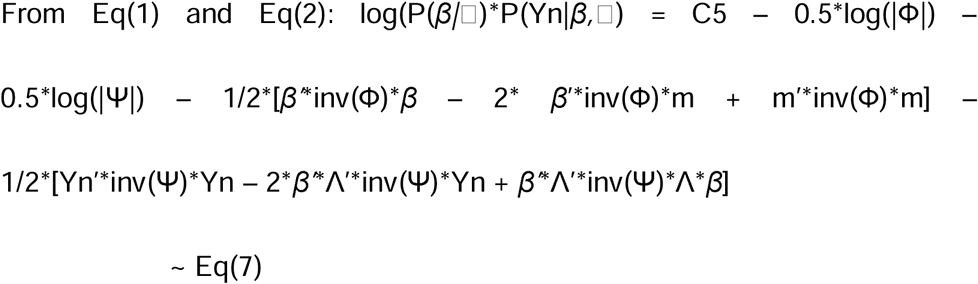

Eq(7) can be divided into 3 parts. Integrating (taking expectation) over Qn(β) for each part with the assistance of Eq(4) and Eq(5) generates:

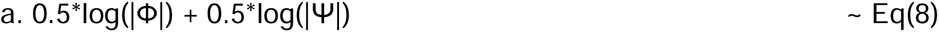

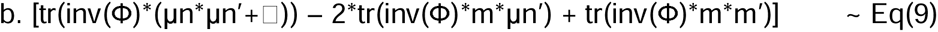

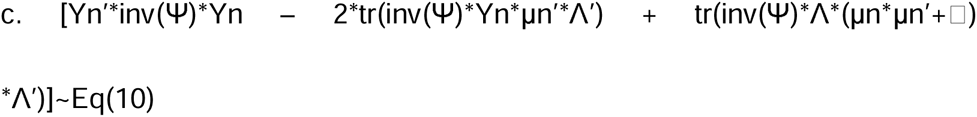

Summation over N samples obtain: F = C – N*Eq(8) – 1/2*∑Eq(9) – 1/2*∑Eq(10).

The covariance matrices Ф and Ψ are both assumed to be symmetric. Now, first, taking derivative of F (or Eq(6)) with respect to **inv(**Ψ) and set it to zero [from Eq(8) and Eq(10)]:

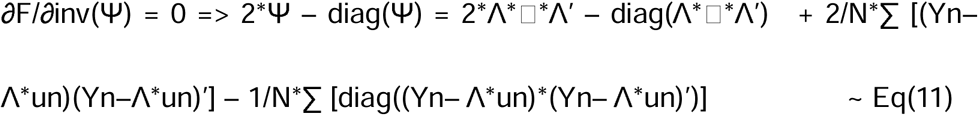

Second, taking derivative of F with respect to **inv(**Ф) and set it to zero [Eq(8) and Eq(9)]:

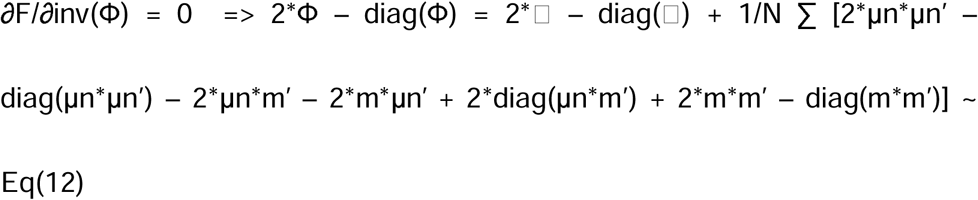

Last, taking derivative of F with respect to **m** and set it to zero [Eq(9)]:

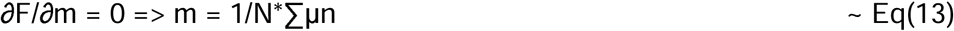

Since μn and □ have been obtained from the preceding E-step, by computing m in Eq(13), Ф in Eq(12) can be determined. Thus, Ψ, m, and Ф (the 3 parameters in set □) that maximize the lower bound of likelihood are all derived.

Note that if Λ is allowed to change with Yn, which will become Λ1, Λ2, Λ3 … coupled with Y1, Y2, Y3 …; and each μn has its correspondent □n, referring to Eq(2) and Eq(3). In this situation, Eq(11) and Eq(12) need to be updated as follows:

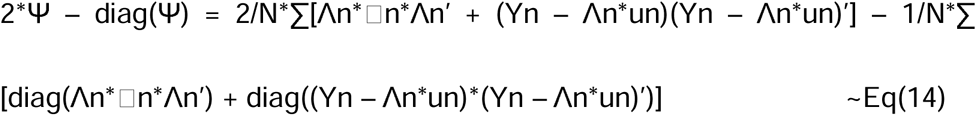

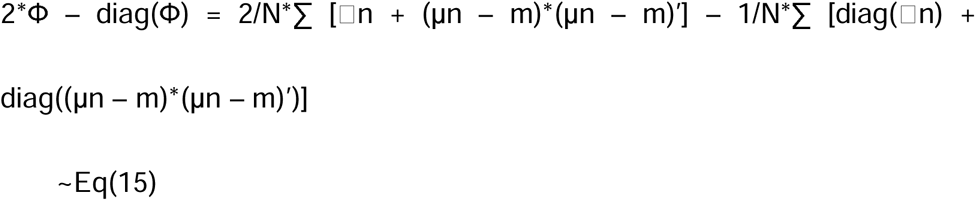

If the covariance matrix was assumed to be diagonal (Ψ or Ф), an extra step of diagonalization would be introduced in the iterations of EM computation.

### Design of simulations to evaluate EMSEV solution performance

The distributions of β are arbitrarily set as follows: mean(β) = [2.5 1.5 0.5 -1] and cov(β)=[2 0.5 0.3 0.2; 0.5 1.5 0.4 0.3; 0.3 0.4 1.8 0.6; 0.2 0.3 0.6 1.7] (cov indicates covariance matrix). The off-diagonal elements indicate dependence between the elements of β. A simpler scenario by setting the off-diagonal elements of cov(β) to zeros was also examined. The mean of ε was zero and its variance was titrated from 1/10 to 10 times of the mean of the diagonal of cov(β), i.e., (2+1.5+1.8+1.7)/4, with intervals of 0.2 (resulting in 11 signal-to-noise ratios in total, SNR). The design matrix Λ was loaded from the built-in sample dataset “airlinesmall.csv” (columns 5 to 8; with Gram-Schmidt orthonormalization) in MATLAB (The MathWorks, Inc., Version R2023a), and simulations were subsequently executed using this software platform.

As for the design matrix, there are specific considerations. Simulations based on EMSEV with a fixed Λ over hundreds of iterations are not inherently problematic but may not be feasible or practical. In most psychological or physiological experiments, subjects undergo examination in only one or a few sessions (which typically comprise hundreds to thousands of trials). In addition, the dimension of β is usually much smaller than that of Y. Consequently, a strategy involving “sliding design matrix” Λn (unit of mini experiment, see **Discussion**), which is a submatrix of Λ, was devised and is detailed below.

In the simulation, the dimension of β is 4×1 and the required sample number is 500. Using a sliding design matrix Λn (4×4) and associated data Yn (4×1)—“sliding strategy I” where Yn, Λn and βn have the same row number—the dimension of Y 2000×1 would suffice (500*4 data points). It is imperative to note that “sample number” N and “data point number” N*dimension-of-Yn are distinct. Since EM iterations can become trapped in local optimum (Dempster et al., 1977), initializing the EM algorithm with a guess approximating the global optimum is highly desirable. A nice feature of sliding design matrix is that it facilitates deriving effective initial estimates of mean and variances (for both β and ε) by resolving the GLM through conventional methods for N times (N = 500 in this case). Strategy I’s limitation emerges when Λn is square and full rank, causing the error term for each ε = Yn – Λn*β to consistently diminish (close to zeros), resulting in suboptimal starting points and increased likelihood of converging to local maxima.

To tackle this issue, an alternative approach termed “sliding strategy II” employed a Yn with vector length of 6×1 and 8×1, representing 50% and 100% increase of the data length in sliding strategy I, respectively. A key advantage of sliding strategy II lies in its ability to provide a suitable starting point while at the expense of incurring 50% to 100% extra data points compared to its predecessor (the sample number or the number of mini experiments remained unchanged). Based on the above observations and inferences, three sets of simulations with different sliding design matrices were explored: (1) dimensions Yn 4×1 and Λn 4×4, and a step size of 4 (Yn to β dimension ratio = 1.0); (2) dimensions Yn 6×1 and Λn 6×4, with a moving step 6 (Yn to β dimension ratio = 1.5); and (3) dimensions Yn 8×1 and Λn 8×4, with a moving step 8 (Yn to β dimension ratio = 2.0). Compared to (2), (3) may provide insights into the potential impact of sample size on the results. Regarding “sliding strategy I”, the initial guesses for cov(β) and var(ε) were set to be identical, specifically one half of the covariance of the estimated β.

Five hundred samples of β and ε, based on pre-defined Gaussian parameters, were generated using the MATLAB function “mvnrnd”. It is noticed that at sample number 500, the covariance of 500 ε samples may not conform to i.i.d. assumption (i.e., the off-diagonal elements are not zeros) and the mean may be different from zeros. To address this issue, the error matrix was centered around the mean, and a transformation guided by Cholesky decomposition was applied, as follows. Assuming the 500 samples of ε generated by “mvnrnd” are denoted by RN_N (already mean-centered and hence, mean(RN_N)=0) with covariance RV (not necessarily diagonal), and the desired covariance matrix is DV (a diagonal matrix). The Cholesky decomposition of RV and DV respectively yields RV_upper and DV_upper. To update RN_N to RN_New with its covariance conforming to DV, the transformation is fulfilled by RN_New = RN_N*inv(RV_upper)*DV_upper. It can be verified that cov(RN_New) = (DV_upper)’* DV_upper = DV. This procedure ensured consistency in the evaluation of simulations.

The same preprocessing was applied to the mvnrnd-generated β samples if their covariance was assumed to be diagonal. With the β and ε samples following their respective desired distributions, Yn can be constructed using the GLM formula. Then, Yn and Λn were input into EMSEV to derive the statistical estimates of β and ε. Meanwhile, in the conventional solution to GLM model, β*n* = inv(Λn’*Λn)* Λn’*Yn and ε*n* = Yn – Λn*β*n*. These conventional solutions (n = 1 to 500) not only serve as initial guesses but also provide another set of distribution for β and ε, which will be compared with the outcomes of EMSEV. The quality of the EMSEV solution was evaluated using the “Frobenius norm relative deviation”, specifically, by computing the Frobenius norm of the difference between the EMSEV-derived and ideal (pre-defined) vector or matrix, divided by the Frobenius norm of the latter. The EM iteration terminated when the difference between the relative deviation ratios of successive iterations fell below 10^-7.

## Results

There are 6 sets of results (full matrix or diagonal matrix of cov(β) x 3 different sliding design matrices), each with 11 relative ratios of var(ε) to cov(β). The pre-defined parameters in distribution were compared with those estimated from EMSEV by relative deviation metrics. The performance of EMSEV was influenced by all the factors addressed in the simulations. It was enhanced by the dimension of Yn (or the number of rows in Λn), the SNR, availability of initial guess provided by the data, and when cov(β) had a simpler structure (i.e., a diagonal covariance matrix). The estimates of the β were quite consistent across all the scenarios. The focus will be on cov(β) and var(ε).

Foremost, the effectiveness of EMSEV is underscored by a significant reduction in percentage deviation of distribution parameters compared to those obtained directly from the GLM over 500 iterations (approximately 6 to 10 times lower for cov(β), and 23 to 29 times lower for var(ε) at SNR = 1; see lower half of Tables 1 and 2). The reliability of the initial guess significantly influenced the performance of the EM algorithm. Results for Λn 4×4 were inferior to those for Λn 6×4 and 8×4, with deviation metrics approximately 1.75 to 2.75 times and 9 to 15 times higher for cov(β) and var(ε) at SNR = 1, respectively (see upper half of Tables 1 and 2). At SNR = 1, the “sliding strategy II” yielded decent results. The deviations for the EMSEV estimates of mean(β), cov(β), and var(ε) were around 3%, 10–16%, and 1.7%, respectively. The detailed results are summarized in Tables 1 and 2, and Figure 1.

**Fig. 1.**
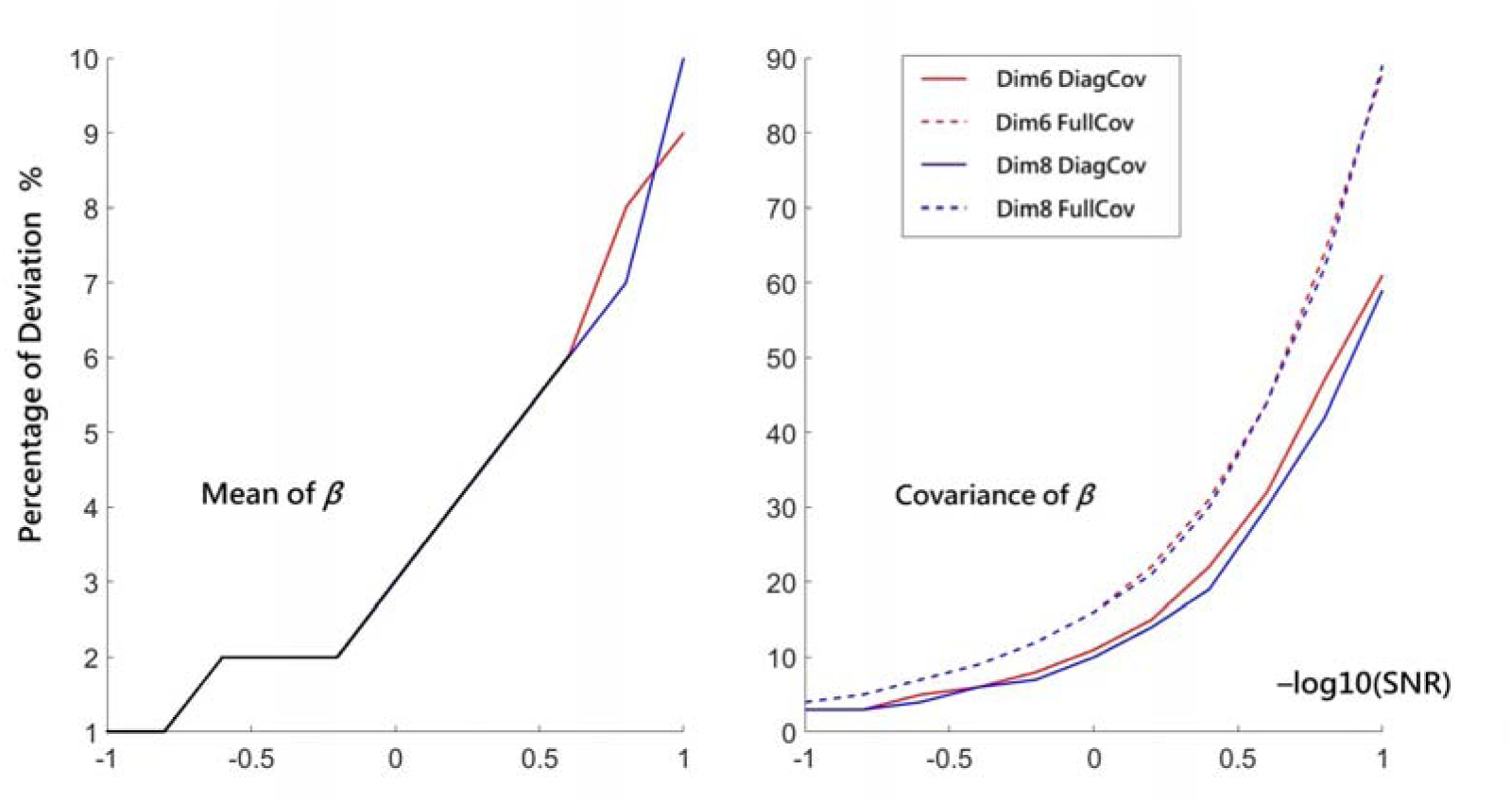
Relationship between signal-to-noise ration (abscissa: SNR, after taking –log10, ranging from 10 to 1/10) and relative deviation from the underlying distribution (ordinate: %). At –log10(SNR) = 0, the variance of β and ε are at the same level. **Left subplot:** Mean of β, with black segments indicating the values of the four lines were very close. **Right subplot:** Covariance of β. **Dim:** dimension of mini experiment matrix Λn (Dim6: 6×4; Dim8: 8×4). **DiagCov:** the covariance of β was diagonalized. **FullCov:** the covariance of β remained unchanged.

**Table 1.** Percentage of relative deviation of 3 distribution parameters for different degrees of signal-to-noise ratio, with default underlying cov(β)

| | dim($\Lambda_n$) | 4x4 | | | 6x4 | | | 8x4 | | |
| --- | --- | --- | --- | --- | --- | --- | --- | --- | --- | --- |
| SNR | $\beta$ | $\text{cov}(\beta)$ | $\text{var}(\varepsilon)$ | | $\beta$ | $\text{cov}(\beta)$ | $\text{var}(\varepsilon)$ | $\beta$ | $\text{cov}(\beta)$ | $\text{var}(\varepsilon)$ |
| 0.1 | 9.39 | 175.42 | 14.27 |  | 9.33 | 87.62 | 2.18 | 9.89 | 88.79 | 1.64 |
| 1.0 | 3.12 | 28.15 | 24.80 |  | 3.01 | 16.06 | 2.89 | 3.07 | 15.88 | 1.75 |
| 10 | 1.02 | 5.15 | 31.32 |  | 0.99 | 4.25 | 2.84 | 0.96 | 4.00 | 1.61 |
| 0.1 | N/A |  |  |  | 9.33 | 927.08 | 66.66 | 9.89 | 931.96 | 50.24 |
| 1.0 |  |  |  |  | 3.01 | 93.53 | 66.91 | 3.07 | 93.49 | 50.21 |
| 10 |  |  |  |  | 0.99 | 10.14 | 66.74 | 0.96 | 10.14 | 49.97 |

Upper half: the outputs of EMSEV; Lower half: the mean and covariance were derived by directly solving GLM 500 times. dim: dimension; cov: covariance.

**Table 2.**
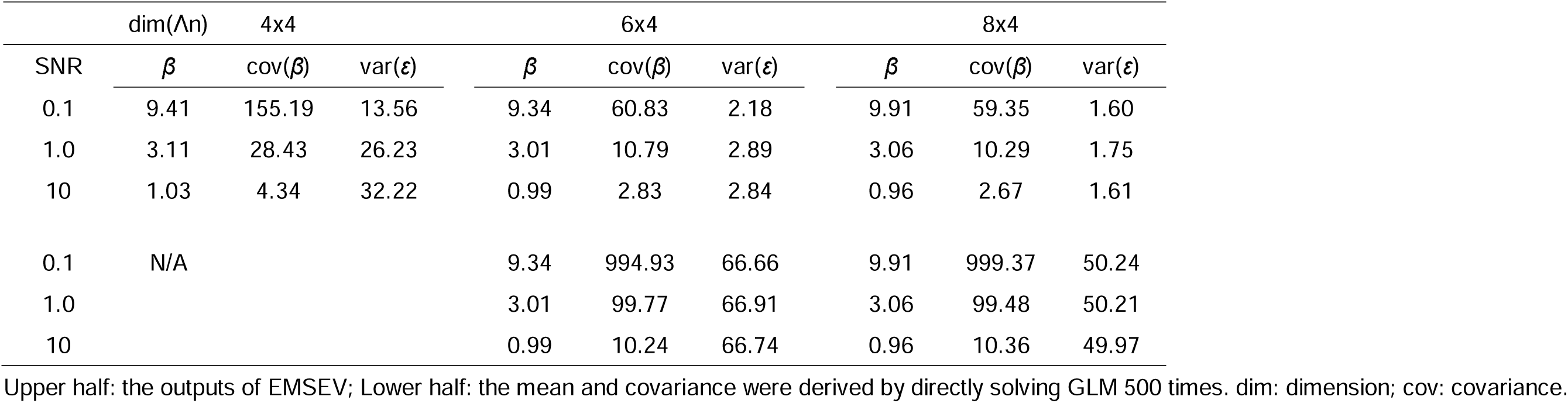
Percentage of relative deviation of 3 distribution parameters for different degrees of signal-to-noise ratio, with the underlying cov(β) diagonalized.

As the noise level increased, both the deviation in the mean and the covariance of β were significantly impacted, with the latter showing a more substantial increase compared to the former. The deviation in var(ε) remained within 1.5 to 3%. At a noise level ten times higher than the signal, the deviation of β reached around 10%, while the deviation of cov(β) could escalate 60 to 90%. This trend is anticipated, as errors in estimating the mean of β amplify the deviation in covariance, thereby accentuating the increase in covariance deviation. Furthermore, in simulations using “sliding strategy II”, where the initial guess was directly derived from the data, the convergence was swift, ranging from seconds to over 10 minutes (depending on the noise level; CPU: Xeon E3-1200 v3/4th Gen Core Processor, Clock Speed: 33 MHz, RAM: 8 GB).

## Discussion

Investigating the variance of biological signals has a longstanding history but has often lacked a robust algorithm capable of distinguishing signal variance from noise (Churchland et al., 2010). This study proposed EMSEV, applying EM algorithm to modified GLM model, as a platform to solve this paramount issue in biological science. The simulation results demonstrate that with appropriate initialization and moderate noise levels, EMSEV shows promise in distinguishing between signal and noise variances. Under the conservative assumption of i.i.d. error terms, the deviation remained below 3% even at SNR 10. When noise and signal levels were comparable, deviations in the mean and covariance of β were approximately 3% and 10–16%, respectively. The ’sliding strategy II’, which incorporates more data points in the mini design matrix and utilizes an initial guess, yielded the best outcomes. It is important to note that the cov(β) and var(ε) obtained through conventional GLM solving were highly biased and are not recommended.

In terms of methodology development, McIntosh et al. introduced a covariance-oriented approach, partial least squares (PLS, invented by econometrician and statistician Wold (Wold, 1966)), to the field of brain science (McIntosh et al., 1996). In their PLS framework, a covariance matrix is constructed from brain imaging data and experimental design (or behavioral profile), which is then decomposed using singular value decomposition. Relevant summary scores indicating the relationship between brain activity and design (or behavior) can be derived, and statistical significance can be assessed using permutation and bootstrap procedures. This technique has been applied to various neuroscientific issues, such as exploring the influence of baseline neural power pattern on event-related activities (Lee et al., 2011). Pascual-Marqui was perhaps the first researcher who treated the variance of signal and noise separately and explicitly in neural models (Pascual-Marqui, 2007). In his formulation of “exact low-resolution brain electromagnetic tomography” (eLORETA), the regularization parameter represents the ratio of measurement noise to biological variance. Both the above canonical methods contribute significantly to the application of (co)variance structures in neuroimaging data. However, they have limitations in effectively quantifying signal variance. By incorporating experiment structure into the design matrix (in contrast to the leadfield of eLORETA that is derived purely from electromagnetic properties), EMSEV provides a novel pipeline aimed at potentially distinguishing between non-biological noise and biological signal variances.

Quantifying the variance of biologically meaningful indices has broad applications. While variability traditionally implied uncertainty (with a negative connotation), recent studies have recognized that variability in biological systems may have “functional” significance. For instance, a certain level of variability in heart rate is linked to better cardiovascular health and increased autonomic flexibility (Thayer et al., 2010). PLS-based analysis has indicated that younger, faster, and more consistent performers exhibit higher brain variability across cognitive tasks such as perceptual matching, attention cueing, and delayed match-to-sample (Garrett et al., 2011), suggesting that increased variability in the central nervous system may support neural efficiency and reduce behavioral variability. Notably, the impact of altering variance is a complex and context-dependent matter. Some research suggests that increased variance in neural signals across specific brain regions may be associated with neuropsychiatric conditions (Scarapicchia et al., 2018). EMSEV could prove beneficial in this relatively new research domain.

In addition, EMSEV offers potential applications to statistical modeling. Unlike current methods that often attribute variance solely to noise under the null hypothesis with a false positive cutoff of p-value 0.05 (where an estimated indicator is compared against a null distribution), EMSEV calculates variances for both target and noise. This enables statistical analysis by comparing two distributions, providing a clear definition of both false positives and negatives (Benjamin et al., 2018). Simultaneous assessment of false positive and negative outcomes in statistics offers practical advantages over traditional methods that focus primarily on false positives. This approach enhances understanding of statistical test performance, enabling researchers to evaluate errors comprehensively and may improve the reliability of findings. Considering both types of errors promotes nuanced interpretation and enhances confidence in statistical conclusions.

To retrieve the distribution of β and ε, each Λn must be of the same size and contain the necessary data to inform all elements in β. The variation is caused by the covariance innate in β and ε. Missing element will bias the estimation of the statistics in Qn(β) and the parameter set □. Therefore, Λn is regarded as a unit of a mini experiment that encompasses all the factors of interest and may represent a miniature of the entire experiment. This appears to be a stringent constraint. The limitation of EMSEV is evident: it operates under the assumption of ergodicity and necessitates substantial regularity in experimental designs. Fortunately, such regularity is common in existing experiments in psychophysics, neurophysiology, neuroimaging, and cognitive psychology. Moreover, if temporal correlation (order of data) is not a concern, the rows of the GLM (observed data and the design matrix) can be permuted and reorganized to ensure that each new Λn carries adequate information of every element of β. The elements of Λ can be continuous or categorical variables, or both. Notably, as demonstrated by the two sliding strategies, it is recommended that the design matrix be constructed to provide a decent initial guess for the EM algorithm. In summary, the total trial number for an experiment may remain constant, but it is suggested that the trails be structured as a mini experiment (Λn). In this mini experiment, all target variables should be present, and the number of data points (rows in Λn) should exceed the number of target variables (columns in Λn).

The “sliding strategy I” yields subpar results compared to its alternatives, which could be partially attributed to the problem of local optima innate in the EM algorithm. To cope with this issue, multiple initializations with diverse starting parameters can be employed, selecting the model with the highest likelihood or lowest loss function. Additionally, heuristic randomization and model refinement techniques have been proposed to enhance the algorithm’s ability to locate global optima (Celeux and Govaert, 1992; Desmond and Glover, 2002). Nevertheless, by applying a sliding design matrix and ensuring that the dimension of Yn is larger than that of β, a well-suited initial guess can be obtained to inform the EM algorithm. It is anticipated that integrating these enhancement strategies into the EM algorithm will further improve the performance of EMSEV, especially in the situation of lower SNR.

## Conclusion

Historically, signal variability has often been attributed to noise in mathematical modeling, despite its acknowledged significance. The application of the EM algorithm within a modified General Linear Model, known as EMSEV, offers a promising approach to estimating both the variances of the target variable and the error. This method holds the potential to effectively address the challenges associated with signal variability and enhance the precision of statistical inferences.

## Authors Contributions

TW Lee is the sole contributor.

## Acknowledgments

This work was supported by NeuroCognitive Institute (NCI) and NCI Clinical Research Foundation Inc. I am grateful for the verification of the formula derivations by Prof. Yu-Te Wu at National Yang Ming Chiao Tung University, Taiwan.

## Financial support

N/A.

## Statements and Declarations

The author declares no conflicts of interest.

## Compliance with ethical standards

No human or animal subject was used in this study.

## Notes

### Competing Interest Statement

The authors have declared no competing interest.

